# Delayed predator response increases ecosystem’s vulnerability to collapse under a changing environment

**DOI:** 10.64898/2026.04.06.716757

**Authors:** Amanda Barreto Campos, Paulo Inácio de Knegt López de Prado, Flávia Maria Darcie Marquitti

**Affiliations:** Programa de Pós-Graduação em Ecologia, Universidade de São Paulo, Instituto de Biociências, São Paulo, Brasil; Universidade de São Paulo, Instituto de Biociências, São Paulo, Brasil; Universidade Estadual de Campinas, Departamento de Genética, Evolução, Microbiologia e Imunologia, Campinas, São Paulo, Brasil

## Abstract

Human activities are driving unprecedented environmental change, yet assessments of ecosystem resilience often overlook the rapid pace of change in the Anthropocene. Predator-prey systems are sensitive to the rate of environmental change and the whole system can collapse if predator population fail to promptly adjust to environmentally-driven shifts in resource population. Here, we investigate how different combinations of predator responsiveness and rates of environmental change influence the system vulnerability to critical transitions, explicitly addressing its interplay with magnitude of change. We found that, as predator responsiveness decreases, relatively slower rates and smaller magnitudes of environmental change leads to system collapse. Hence, even low and seemingly inoffensive total magnitudes of environmental change can be catastrophic if the rate of change is beyond a critical threshold. We propose considering predator responsiveness and current rates of environmental change as crucial factors in predicting the Anthropocene’s impact on ecosystems.

## Introduction

Human activities are causing unprecedented environmental changes across ecosystems (Joos and Spahni, 2008). Increasing mean temperature, pollution, habitat loss and fragmentation have affected all levels of biological organization (Joos and Spahni, 2008; Klein Goldewijk et al., 2011; Hughes et al., 2017; Zanette et al., 2000; Fischer and Lindenmayer, 2007). As external conditions change, ecological systems can suddenly shift to a qualitatively different regime or state. This phenomenon is known as critical transition (Scheffer and Carpenter, 2003) and describe events such as rain forest savannization (Hirota et al., 2011) and coral reef depauperation (Hughes et al., 2017).

Critical transitions are often approached through the lenses of steady state analysis of dynamic models, where shifts are expressed as changes in the basin of attraction of the stationary states (Ashwin et al., 2012; Scheffer and Carpenter, 2003; Scheffer, 2020; Holling, 1973). The goal of this approach is to determine the magnitude of external environmental change or disturbance a system can withstand until it shifts to another state to gauge its vulnerability and resilience (Scheffer et al., 2001; Holling, 1973). Though this approach allowed important advances in our understanding of system vulnerability under static or slowly changing environmental conditions, recent theoretical research put in check if it can properly describe the observed pace of environmental change of the Anthropocene (Donohue et al., 2016; Siteur et al., 2016; Pinek et al., 2020; Synodinos et al., 2023; Vanselow et al., 2019).

Theoretical work explicitly addressing the rate of environmental change found that critical transitions can occur in the absence of changes in the basin of attraction of the ideal system state, that is, with no apparent loss of stability (Ashwin et al., 2012; Siteur et al., 2016; Vanselow et al., 2019). When the environmental conditions changes at a rate beyond the system’s ability to keep pace of the shifting attractor, the ecosystem suffers a qualitative change, leading to what is known as a rate-induced transition (Synodinos et al., 2023; Siteur et al., 2016; Ashwin et al., 2012). In fact, system transitions can occur earlier than predicted solely by steady state analysis when the rate of environmental change is also considered (Ashwin et al., 2012). In this sense, our expectation of resilience of a plethora of ecosystems are potentially overestimated.

Recent studies found that predator-prey interactions are rate-sensitive. Theory suggest that when environmental conditions force prey growth rate (Siteur et al., 2016; Vanselow et al., 2022; Kaur and Dutta, 2022) or prey intraspecific competition (Vanselow et al., 2019) to change above a critical rate the whole ecosystem collapses. The collapse occur because changes in predator population abundance are slower than environmentally-driven changes in the prey population abundance (Vanselow et al., 2019; Siteur et al., 2016). Although the pace at which the predator population respond to changing prey population - predator responsiveness - is one of the key elements determining system collapse (Siteur et al., 2016), all previous studies considered that the predator population have a fixed capacity to respond to changes in prey population.

The responsiveness of predators population can derive from the resultant difference of populational characteristic times of predator and prey. Empirically, predators tend to have larger body sizes than their preys (Cohen et al., 1993) and thus longer lifespans. In this sense, predator population can have a slower charateristic time than its prey. Since the average predator-prey body-size ratio differ across habitats (Brose et al., 2006), the consideration of different predator’s responsiveness to environmentally-driven prey population change can give insightful information on the vulnerability of different ecosystems to collapse.

In the present work, we used a non-autonomous Rosenzweig-MacArthur predator-prey model with timescale separation between populations to investigate system vulnerability to critical transitions. We found that the vulnerability in this system to undergo rate-induced transition depends on the consumer responsiveness and the rate in which environmental degradation increases. Rate-induced transitions adds another layer of complexity on our understanding of ecosystem resilience, previously focused on the amount of disturbance and environmental change a system can withstand before shifting to a qualitatively different regime.

## Material and Methods

### The slow-fast Rosenzweig-MacArthur predator-prey system

We used a time-scaled Rosenzweig-MacArthur model (Rosenzweig and MacArthur, 1963) with rescaled prey (*R*) and predator (*C*) populations relative to carrying capacity (Eq. System 0.1).

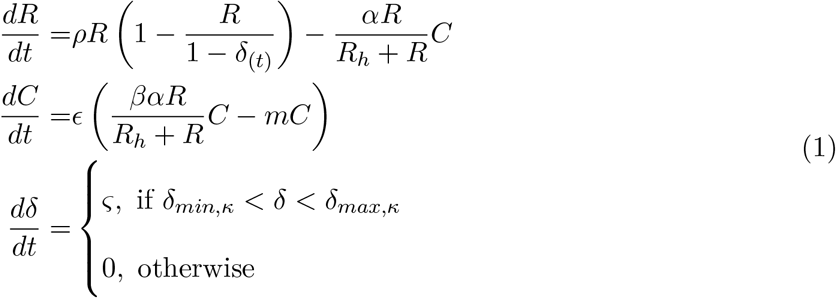

Briefly, the prey population increases with intrinsic growth rate (*ρ*) until it saturates (1 − *δ*), reaching its maximum population size in the absence of environmental degradation (*δ* = 0). Prey population decreases with predation by predators which have an attack rate (*α*) and a functional response with a half-saturation constant (*R*_*h*_). Predators increase in population as predated prey are converted into new predator individuals in a efficiency proportion constant (*β*) and decrease as they decay in a constant death rate (*m*).

We explicitly model two key parameters: the responsiveness of predator population (*ϵ*) and the environmental degradation (*δ*). The responsiveness of predator population (*ϵ*) represents a separation in the characteristic times of predator and prey population (Rinaldi and Scheffer, 2000; Siteur et al., 2016; Vanselow et al., 2019). As *ϵ* tends to zero, predator population responsiveness to changes in prey population is slower. The effect of environmental degradation (*δ*) is explicitly addressed as a process decreasing the saturation point of prey population. The environmental degradation can be constant 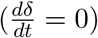 or increase over time in a rate *ς* within a bounded magnitude of environmental degradation (*δ*) (see Appendix A for details). Once the rate of environmental change reaches its maximum value (*δ* = *δ*_*max*_) it becomes constant over time.

### Model parametrization and definition of system collapse

Here we are interested in the emerging patterns of predator-prey systems under different scenarios of changing environment. The model is parameterized in a way that the phase space is permanently limited to a region where predator and prey coexist in an stable equilibrium point (see Supp. Table 1 for the comprehensive list of parameter values).

We used LSODA from scipy.integrate package to approximate the numerical solution for each combination of consumer responsiveness (*ϵ*) and rate of environmental degradation (*ς*). Since fast-slow systems are known to have “stiff equations” (Rinaldi and Scheffer, 2000) we log-transformed the equations and used strict values for the integration parameters *hmax* = 0.01 and *mxstep* = 10000 to improve the quality of numerical approximations. We initialized all simulations considering the initial prey and predator population equivalent to the values they assume at the coexistence equilibrium point in absence of environmental degradation (*δ* = 0).

The prey population is re-scaled relative to its carrying capacity, hence the maximum prey population is 1. Here we defined as system collapse the case where prey population crosses the extinction threshold of 10^−19^. We consider that the prey population is virtually extincted for this threshold since a small demographic perturbation or noise could lead to local extinction (Liebhold and Bascompte, 2003).

## Results

### The magnitude of environmental degradation decreases predator’s population

The fast-slow Rosenzweig-MacArthur (RM) is qualitatively equivalent to the classic RM model in terms of stationary states. In the absence of rate of environmental degradation 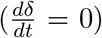,both models have the same three stationary points: Extinction (*e*_1_), Prey Release (*e*_2_) and Predator-Prey Coexistence (*e*_3_). For both models, the magnitude of environmental degradation (*δ*) but not the predator responsiveness (*ϵ*) influence the value that predator populations assume on the predator-prey coexistence (*e*_3_) equilibrium (Fig. 1 and Supp. Fig. 5). However, in our fast-slow RM model we do not consider the case where predator and prey coexist in a cyclic dynamics, since we bounded our system to a maximum *δ* numerical value below the threshold value for such cycles. Hence, we have not included the classical case of the enrichment paradox (Rosenzweig, 1971) in our study.

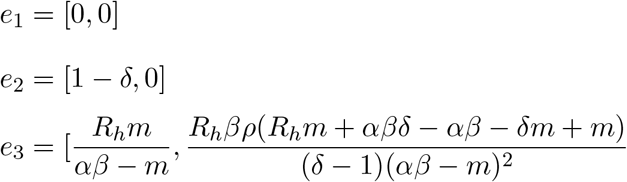

**Figure 1:**
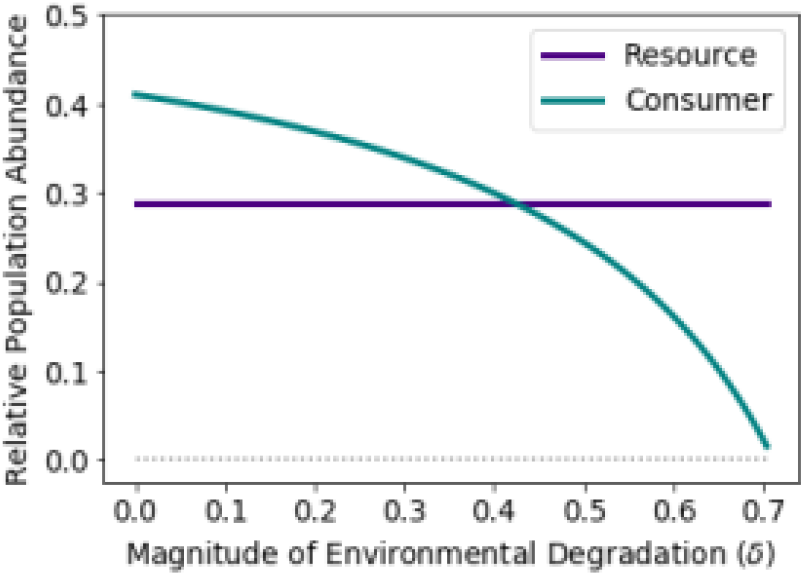
The effect of the *magnitude* of environmental degradation on population abundance at equilibrium. We bounded our analysis within values of magnitude of environmental degradation (*δ*) that corresponds to the predator and prey coexistence in a equilibrium point. By using the coexistence equilibrium expression *e*_3_ for prey and predator we found that, as environmental degradation (*δ*) increases, the predator population at equilibrium decreases while the prey population remains constant.

To investigate the effect of the rate of change 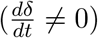 on the coexistence equilibrium of predator and prey (*e*_3_), we restricted the phase space of our model to the corresponding region by restricting the values of *δ* (parametrization details in Supp. Information 0.2). In this sense we guarantee that the asymptotic behaviour of the system is where both populations abundances are larger than zero. Therefore, differently from the classic RM model, our working model has a single possible equilibrium: the coexistence of predators and preys. There is no possibility of bifurcations for the chosen parameter space in the absence of a rate of environmental change.

The population of predators on equilibrium decreases as the magnitude of environmental degradation increases, hence the predator population have its maximum population size in a pristine environment (*δ*=0), and its minimal population when the magnitude of environmental degradation is maximal. Note that the prey population on equilibrium remains constant as the magnitude of environmental degradation increases (Fig. 1). We will investigate the effect of the rate of environmental change 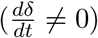 considering this restricted phase space.

### The rate of environmental degradation leads to prey population collapse

We relaxed the assumption that environmental degradation (*δ*) remains constant until the system reaches its stable equilibrium made in the previous section to study the effect of the rate of environmental degradation on population abundance 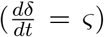.We begin all simulations with initial populations values that prey and predator assume at a pristine environment (*δ* = 0). Considering a fixed predator responsiveness, we set the system to reach the same maximum magnitude of environmental degradation at different rates. The system takes dramatically distinct trajectories for slightly different rates of environmental degradation (Fig. 2). When the rate of environmental degradation reaches a critical value, the prey population cross the extinction threshold and the system collapses (Fig. 2A,C). We consider it a critical rate of environmental degradation 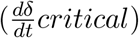 for this system. Hence, when the environmental degradation is slower than the critical rate, the prey population never cross the extinction threshold (Fig. 2A,B).

**Figure 2:**
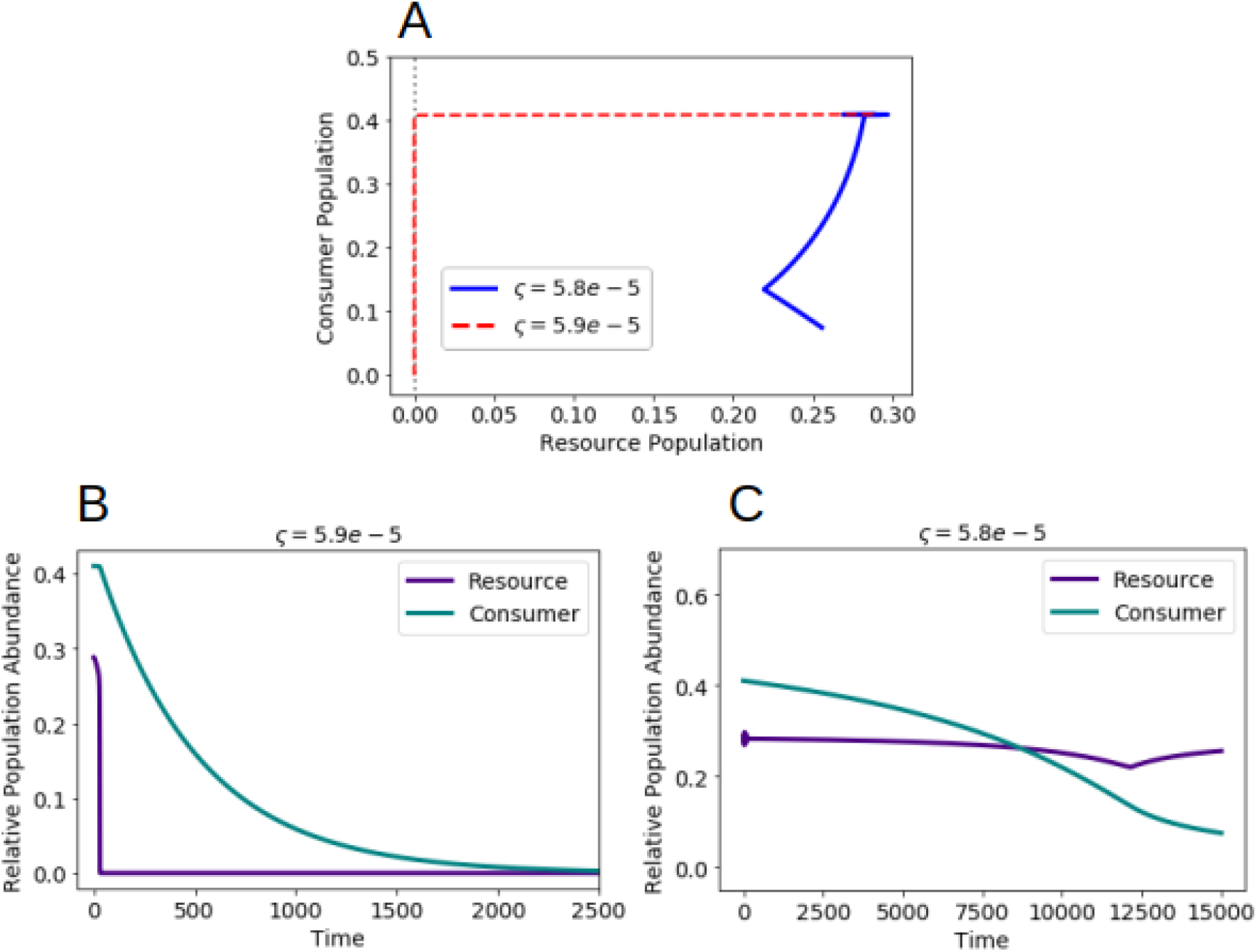
The effect of the *rate* of environmental degradation on population abundance at equilibrium. (A) For slightly different rates of environmental change 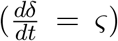 the system take distinct trajectories highlighted in the phase space portrait. (B) When the environmental degradation reaches a critical rate 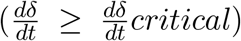, the prey population cross the extinction threshold and the system collapses. (C) When the environmental degradation is slower than the critical rate 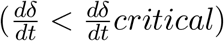, the prey population never cross the extinction threshold.

When we assume that the environmental degradation remains constant 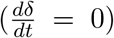 until the system reaches its stable equilibrium or that it changes at a given rate bellow a critical value 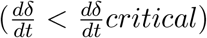, the system responds timely to the environmental degradation (Fig. 1B, Fig. 2C respectively). However, when the rate of environmental degradation crosses a critical value 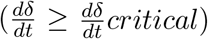 the system is unable to track the moving equilibrium leading to collapse (Fig. 2B). Note that, when we disregard the effect of the rate of environmental degradation, the predators population abundance, not the prey’ reduce in front of increasing magnitude of environmental degradation (Fig. 1 and equilibrium expression *e*_3_). However, when we consider the rate of environmental degradation the prey population crashes prior predator’s population (Fig. 2B).

### Slow predator responsiveness increases the system vulnerability to rate-induced collapse

We investigated the interplay between the different degrees of responsiveness of predator population to changes in prey population (*ϵ*) and different rates of environmental degradation 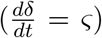 . We consider that each value of *ϵ* represents a combination of predator-prey timescale separation, therefore a different consumer-resource system. The rates of environmental degradation 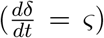 are external factors driving system change, hence they influence but are not influenced by the system. Each rate represents a scenario of environmental degradation.

The vulnerability of the system to undergo rate-induced collapse is determined by the interplay between the intrinsic responsiveness of predator and the rate of environmental degradation. We found that for the majority of predator responsiveness values (*ϵ*) there is a threshold of rate of environmental degradation 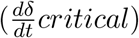 for which the system collapses (transition from the grey to the colored area of the Fig. 3). When the rate environmental degradation is bellow a critical threshold 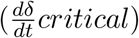 for a given *ϵ* there is no system collapse (grey area of the Fig. 3). Note that as the predator responsiveness decreases (*ϵ* → 0.001) the critical rate environmental degradation for which the system collapse 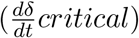becomes smaller. Hence, systems where the predator population reacts slower to changes in the resource population are more vulnerable to rate-induced collapse.

**Figure 3:**
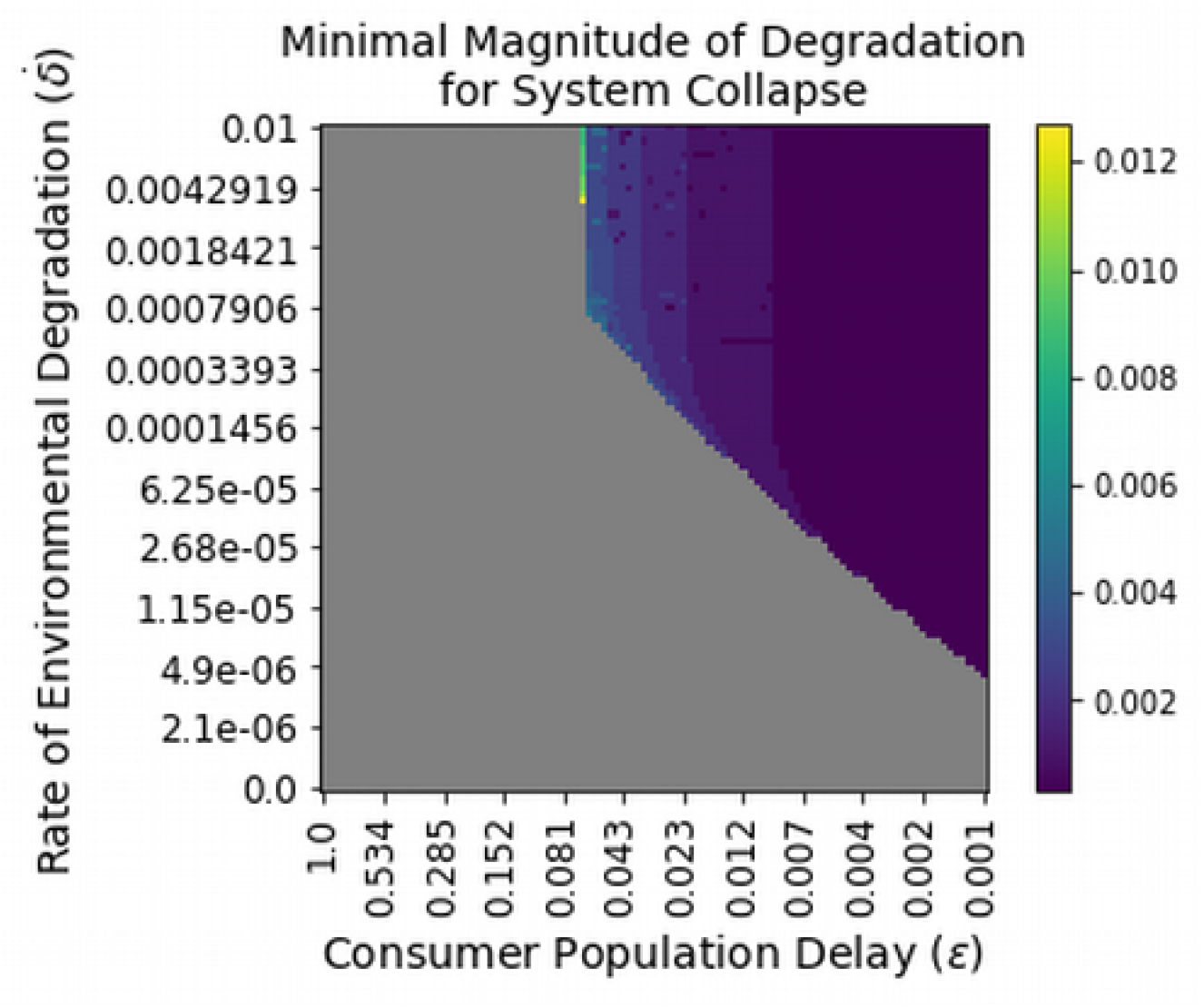
Effect of consumer responsiveness on the system’s vulnerability to rate-induced collapse. The color grey is the absence of rate-induced collapse, whereas the colored area represent system collapse (*R <* 10^−19^). The color bar represents the critical magnitude of change (*δ*_*critical*_) for which the system starts to be vulnerable to rate-induced collapse. We set all simulations with initial populations values that resource and consumer assume in a pristine environment (*δ* = 0). We included the particular case where 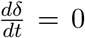 to demonstrate that in the absence of a *rate* of environmental degradation, the system does not collapse regardless of consumer responsiveness (*ϵ*). For the majority of consumer responsiveness (*ϵ*) there is a threshold of rate of environmental degradation 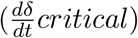 that, when crossed, the system collapses.

### Predator’s responsiveness determines the critical magnitude of environmental degradation

Considering every combination of rate of environmental degradation 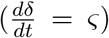 and predator responsiveness (*ϵ*), we determined the critical magnitude of change for which the system becomes susceptible to rate-induced collapse (*δ*_*critical*_), depicted in the color bar of Fig. 3. We show that *δ*_*critical*_ values represent up to 2% of the total potential magnitude of environmental degradation (*δ*_*max*_ = 0.704) (Fig. 3). Since this percentage is low, we can imply that the system is virtually extincted for any magnitude of environmental degradation when the system reaches the critical rate of environmental degradation 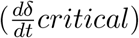.

We found that for slightly different magnitudes of environmental degradation (*δ*_*max*_), the prey population either cross or not the extinction threshold (Fig. 4A-B). Predator responsiveness (*ϵ*) influences the value of critical magnitude of environmental degradation for which the system is prone to collapse (Fig. 4C). We show that, as the predator population responds faster to the changes in prey population (*ϵ* → 1) the magnitude of environmental degradation for which the system begins to be susceptible to rate-induced collapse (*δ*_*critical*_) increases (Fig. 4C). That implies that the system becomes relatively more resilient to rate-induced collapse as the predator is able to respond timely to changes in prey population.

**Figure 4:**
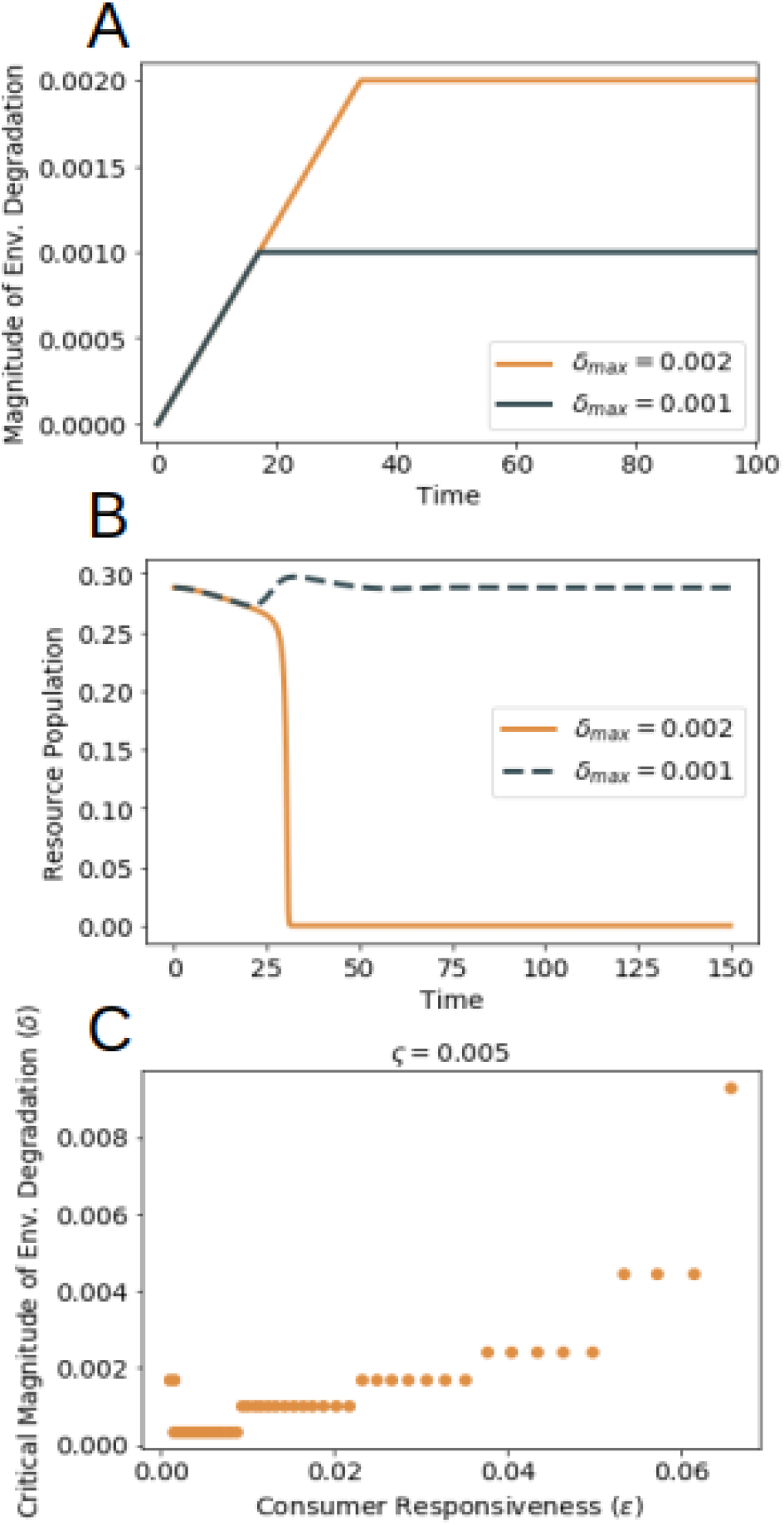
Critical magnitude of environmental degradation 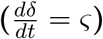 determine system’s susceptibility to rate-induced collapse. Considering the same rate of environmental degradation 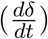and consumer responsiveness (*ϵ*), we tested how different maximal magnitudes of change (*δ*_*max*_) influence system’s susceptibility to rate-induced collapse. (A) For slightly different magnitudes of change (*δ*_*max*_), (B) the resource population either cross or not the extinction threshold. When the resource population crosses the extinction threshold the *δ*_*max*_ is considered as a critical magnitude of change (*δ*_*critical*_). (C) Note that as consumer responsiveness to changes in the resource population improves, the value of critical magnitude of change (*δ*_*critical*_) increases. That implies that the system becomes relatively more resilient to rate-induced collapse.

## Discussion

In this work we moved beyond a qualitative approach of fast/slow rate and large/small magnitude of environmental change and uncovered a general pattern of system vulnerability to rate-induced collapse. We demonstrated how these theoretical qualifications are system-dependent by explicitly investigating the interplay between rate of environmental change, magnitude of environmental change and system responsiveness.

Predator responsiveness does not influence the abundance of either predator or prey population on equilibrium, however our results show that it is a key property determining ecosystems’ vulnerability to collapse induced by environmental change. In our model, predator responsiveness is the timescale difference between prey and predator characteristic times. Such difference can occur because predators usually have larger body sizes than its prey (Cohen et al., 1993; Brose et al., 2006), which can be related to longer lifespans and longer population turnover. We found that, as predator responsiveness decreases, relatively slower rates and smaller magnitudes of environmental change lead to system collapse. Hence, our results suggest that ecological systems where the predator population is less responsive to changes in the prey population are more vulnerable to rate-induced collapse. In freshwater habitats, predators have, on average, larger body sizes than their prey in comparison to marine and terrestrial habitats. Additionally, the body-size ratio of vertebrate predators and their prey usually are higher than the body-size ratio of invertebrates predators (Brose et al., 2006). From these pattern we can imply that freshwater vertebrate predator-prey systems could be more vulnerable to rate-induced collapse due to the potentially larger difference in characteristic times between predator and prey in comparison to invertebrate predator-prey marine and terrestrial ecosystems. Our results demonstrate that the critical threshold of environmental degradation for either rate or magnitude is contingent on evolutionary history and ecological features of each system. Therefore, our modeling approach shows an instance by which species traits affect the dynamic of interactions and the system vulnerability.

By considering the fundamental unit of trophic networks - the pairwise relationship between prey and predator - we investigated the case that predators are specialized in a single prey. In this system, collapse occurs due to the predator over-consuming the prey (Siteur et al., 2016; Vanselow et al., 2019). Here we considered that environmental degradation decreases prey population sizes only. Consequently, specialized predators populations are indirectly affected by the environmental degradation. In fact, diet specialization is related to greater extinction risk under changing environmental conditions (Colles et al., 2009). A step forward is to address how the rates of environmental degradation could modulate complex predator-prey interaction. There is theoretical evidence suggesting that the magnitude of environmental change can destabilize multi-species trophic networks (Kaur and Dutta, 2020), however, its vulnerability to rate-induced collapse is still to be assessed.

We considered environmental change as a process (chronic disturbance) rather than an event (acute disturbance) (Pinek et al., 2020). Particularly, we studied an environmental disturbance gradually reducing prey carrying capacity. Since prey carrying capacity is related to habitat use, disturbances impacting the habitat influence the maximum size of the prey population. This process is both specific enough to provide a straightforward ecological interpretation and broad enough to encompass various causes. For instance, in terrestrial systems, environmental disturbances can be caused by habitat loss, degradation and fragmentation (Zanette et al., 2000; Fischer and Linden-mayer, 2007; Klein Goldewijk et al., 2011). The rate of environmental change considered here has the same general behaviour as the chronic press disturbance assessed in conservation biology: an increasing disturbance (ramp stage) followed by a period where the disturbance is constant (stasis stage) (Pinek et al., 2020), which is sustained for a relevant period of time. Here we explicitly addressed how the ramp stage of the disturbance affects a predator-prey system. We show that the consequences of press disturbances on ecosystems are not solely determined by their total magnitude but are primarily influenced by the speed of the ramp stage. Our results demonstrate that once the environmental change surpasses a critical rate (ramp stage angle), the maximal magnitude of change the system can withstand without collapsing is negligible. Therefore, even seemingly inoffensive total magnitudes of press disturbances can have catastrophic consequences if the ramp stage exceeds a critical threshold. It’s noteworthy that both chronic press disturbances and acute pulse disturbances exhibit a ramp stage. Hence we hypothesize that rate sensitivity may be an overlooked characteristic of both types of environmental disturbance.

Most of disturbances in nature are well characterized by changes in the magnitude of disturbance over time (Donohue et al., 2016). In fact, there are theoretical, observational and experimental evidences suggesting the rate-sensitivity of systems across all biological organizations levels (Pinek et al., 2020). For instance, an experimental study found that soil mycorrhizal community structure changes can be sensitive to the rate of increasing *CO*_2_ concentrations, where the same magnitude of change had no significant effect (Klironomos et al., 2005). Even though the relevance of rate of change have been empirically addressed, few experimental studies provided detailed information on the magnitude of change and the time passed during the ramp stage (Pinek et al., 2020). The elaboration of a clear and precise reporting of this information would facilitate comparison across studies, hence emphasizing the value of having a unified framework for designing and reporting such experiments (Pinek et al., 2020). Generally, empirical studies face an additional challenge to untangle the effect of the magnitude, the rate of change and the duration of the disturbance, which is necessary to correctly infer the rate-sensitivity of the system (Synodinos et al., 2023; Pinek et al., 2020).

Historically, resilience has been conceptually defined as the capacity of the system recover from a disturbance (Scheffer, 2020). Hence, a larger resilience is equivalent to a faster recovery. This concept is often operationally assessed in terms of dynamical system theory as how large and negative is the real part of the eigenvalue of the stationary point of interest. From this we determine the stability of a given system state, which is translated to the size of the attracting basin (Scheffer, 2020). A transition would occur once a disturbance is large enough to cause the system to escape from its current attracting basin and diverge to another state (Holling, 1973; Scheffer, 2020). Here we show that system’s vulnerability to collapse increases with the rate of environmental degradation, even in the absence of changes in the eigenvalues and basin of attraction of the ideal system state (Siteur et al., 2016; Vanselow et al., 2019; Synodinos et al., 2023). The system collapses due to the dramatic decrease in prey population density during the transient trajectory, rendering it highly vulnerable to extinction. Transients has been already recognized as relevant to understand ecological systems, however the focus has been on how long the system remains from its asymptotic behaviour mimicking an apparent regime shift (Hastings et al., 2018). In our work we show that how far the system gets from the asymptotic behaviour before converging can also be a relevant aspect of the transient since the population abundance gets so small that a dramatic transition can occur in the system due to stochastic events and environmental noise (Liebhold and Bascompte, 2003). In this sense, we recommend ecological resilience to be revisited to include information on the transient of the system.

The increasing collection of experimental, observational and theoretical evidence supporting the significance of rate-induced collapse for ecological systems (Synodinos et al., 2023) alongside our results endorse that the current rates of change could be an equally or even a more important aspect to evaluate in global change scenarios projections that currently rely primarily on the total magnitude of change for a given time frame (Steffen et al., 2015). Here we provide evidence supporting the inclusion of the response potential of the system under changing environment on an ecological level as an important aspect to consider when assessing the impact of the Anthropocene on ecosystems. We endorse that solely considering the magnitude of environmental change, and overlooking the rate of change, can ill-inform our management decisions and conservation efforts jeopardizing the future of ecosystems (Joos and Spahni, 2008)

## Acknowledgements

This study was financed in part by the Coordenação de Aperfeiçoamento de Pessoal de Nível Superior – Brasil (CAPES) – Finance Code 001 – FMDM). ABC was funded by grant #2022/06847-4, São Paulo Research Foundation (FAPESP). The authors thank Rafael Menezes and Renato Coutinho for assistance with computational challenges, Guilherme Longo, Lisa McManus, Marina Hirota, and Vitor Vasconcelos for insightful comments on the earlier drafts of the manuscript.

## Supplementary Information

### 0.1 Re-scaling Rosenzweig-MacArthur consumer-resource model

The fast-slow predator-prey model is defined as follows (Eq. System 2):

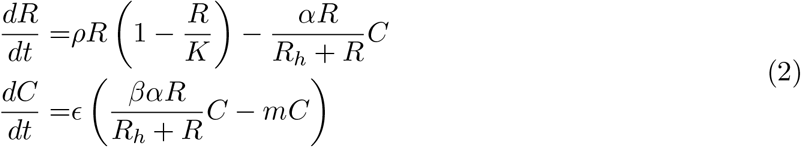

Briefly, the prey population increases with intrinsic growth rate (*ρ*) until it saturates (*K*). The prey population decreases with predation by predators (*C*) which have an attack rate (*α*) and a half-saturation constant (*R*_*h*_) until it saturates. Predators increases in population as predated resources are converted into new predator individuals in a efficiency proportion constant (*β*) and decrease as they decay in a constant rate (*m*). For the particular parameter value *ϵ*=1 we recover the original model Rosenzweig and MacArthur (1963). For this model there are four equilibria: extinction equilibrium point (*e*_1_), prey release equilibrium point (*e*_2_), predator-prey coexistence in a equilibrium point (*e*_3_) and predator-prey coexistence in a limit cycle (Supp. Fig. 5).

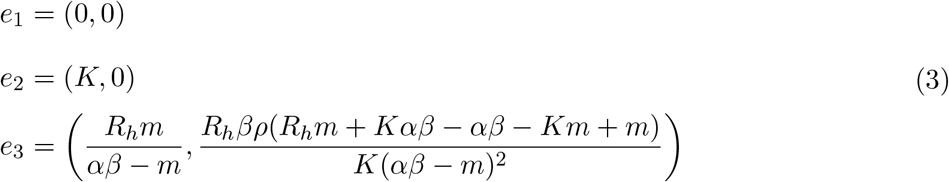

Since we wanted to analyse the effect of rate of environmental change, we delimited the parameter space of the system so no bifurcation would occur. First, we re-defined the carrying capacity (*K*) to explicitly consider the environmental degradation relative to the carrying capacity, therefore *K* = *K*_*max*_ − *δ*. The *K*_*max*_ is the value of *K* in which the system transitions from the coexistence in equilibrium point to a coexistence in a limit cycle. Since we wanted to analyse the system exclusively on the coexistence equilibrium point, we set this threshold as the maximum value of *K* and re-scaled the system. Hence this model has no enrichment paradox (Rosenzweig, 1971). Additionally, we re-scaled the prey and predator populations in terms of units of carrying capacity (*K*). To do that we re-scaled *R, C, R*_*h*_ and *δ* in terms of *K*_*max*_. Finally, we added a new equation to explicitly address how the environmental degradation (*δ*) changes over time (*ς*). The working model is (Eq. System 4):

**Figure 5:**
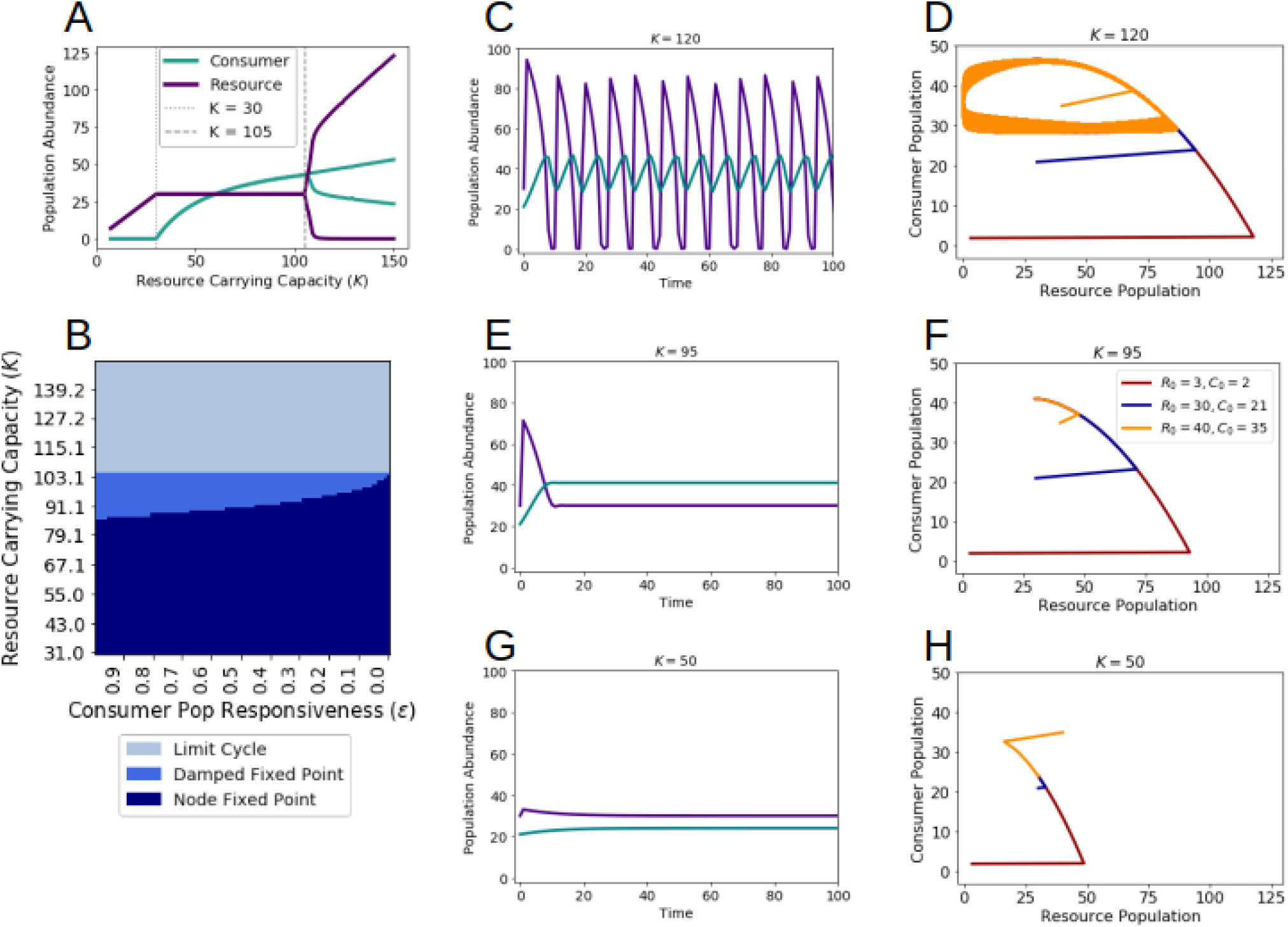
Different regimes of the fast-slow predator-prey system. The fast-slow Rosenzweig-MacArthur predator-prey model have three asymptotic non-trivial regimes. (A) Prey release (*K <* 30), Coexistence in a stationary equilibrium (30 *< K <* 105) and Coexistence in a limit cycle (105 *< K <* 150). Note that these threshold values are associated to the parameter space considered in this work. (B) Since we are interested solely on the coexistence of predators and prey, we calculated the trace and determinant of the Jacobian matrix for this equilibrium and identified three regimes: node fixed point, damped oscillations fixed point, limit cycle. (C) temporal evolution and (D) phase space of a limit cycle. (E) temporal evolution and (F) phase space of damped oscillations fixed point. (E) temporal evolution and (F) phase space of a node fixed point. Each line in the phase space (D, F, H) represents the trajectory of the system from a different initial value of predator and prey.

**Table 1:** Supp. Table 1. Comprehensive parameter list.

| Symbol | Parameter | Value |
| --- | --- | --- |
| $\rho$ | prey intrinsic growth rate | 40 |
| $\delta$ | magnitude of environmental degradation | [0, 0.704] |
| $R_h$ | consumption half-saturation | 0.432 |
| $\alpha$ | predator maximum attack rate | 50 |
| $\beta$ | predator conversion efficiency | 0.01 |
| $m$ | predator mortality | 0.2 |
| $\epsilon$ | predator responsiveness | [0.001, 1] |
| $\varsigma$ | rate of of environmental degradation | [0, 0.01] |
| $R_o$ | initial prey population | 0.288 |
| $C_o$ | initial predator population | 0.410112 |
| $\delta_o$ | initial magnitude of environmental degradation | 0 |
| $x$ | prey extinction threshold | $10^{19}$ |

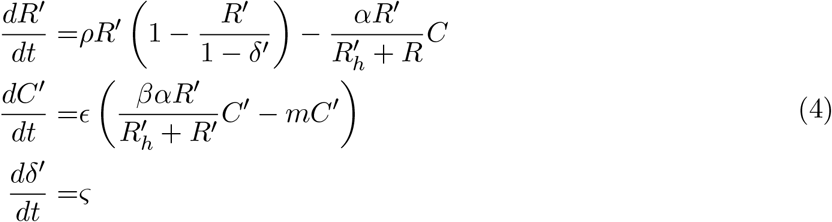

### 0.2 Model Parametrization

We chose model parameter values to permanently limit the phase space to a region where predator and prey population coexist in a equilibrium point attractor to allow us investigate solely the effect of the different rates of environmental degradation on the ecosystem (Supp. Table 1). We explicitly model two key parameters: the responsiveness of predator population (*ϵ*) and the rate of environmental degradation (*ς*). The responsiveness of predator population (*ϵ*) represents a time scale separation between prey and predator characteristic times to represent their differences in population turnover (Rinaldi and Scheffer, 2000), and as *ϵ* tends to zero, predator population responsiveness to changes in prey population is slower. The environmental degradation (*δ*) is explicitly addressed as a process decreasing the saturation point of prey population. The environmental degradation increase over time in a constant rate *ς* until it reaches the maximum value (*δ* = *δ*_*max*_) and becomes constant over time.

### 0.3 Implemented Mathematical Model

We log-transformed the model to cope with integration challenges of stiff equations. Consequently, in this model the population values approach but never reach zero. The differential equation system used to integrate the system is the following (Eq. System 5):

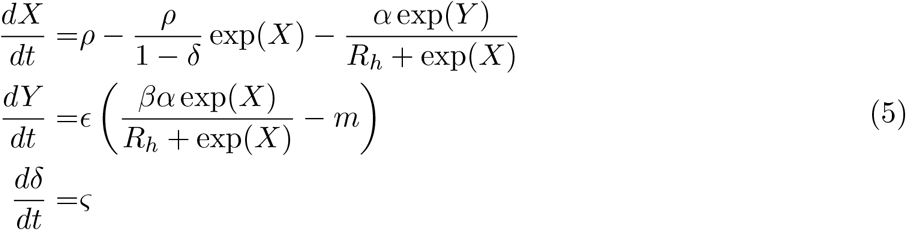

We initialized the system with log transformed predator and prey populations values (*X* = log(*R*) and *Y* = *log*(*C*)) corresponding to the values they assume in a pristine environment (*δ* = 0). We took the exp from the resultant population values and used them on the subsequent analysis.

### 0.4 Rate-Induced Collapse is not a classical Critical Transition

We consider that the resource population is virtually extincted when it crosses the threshold of 10^−19^ since a small demographic perturbation or noise could lead to local extinction (Liebhold and Bascompte, 2003).However, since there is no change in the stability of the equilibrium, the system eventually converges to the equilibrium value expected analytically (Supp. Fig. 6). Bellow there are the system trajectories without the extinction threshold (Supp. Fig. 7).

**Figure 6:**
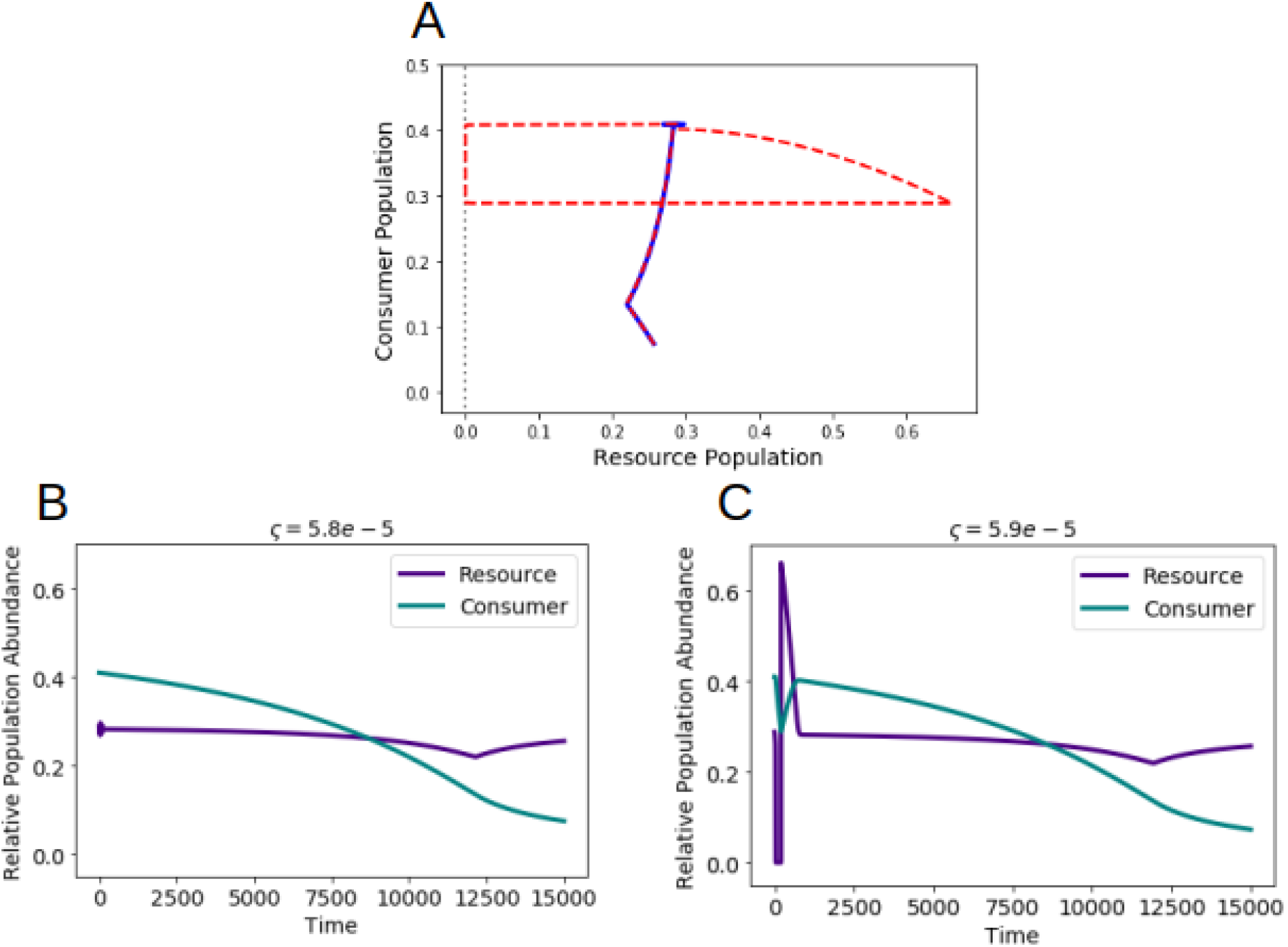
Effect of rate of environmental degradation on population abundance at equilibrium. (A) For very slow and slightly different rates of environmental change 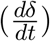 the system take remarkably distinct trajectories. For a slightly faster rate of environmental change (dashed red line) the system reaches the extinction threshold of *R <* 10^−19^. After the prey population reaches this critically low abundance the population increases. This happens because there is no loss in stability of the coexistence fixed point, hence, the system eventually converges to the equilibrium where both population exists. (B) When the the rate of environmental change is slower, the prey population never cross the extinction threshold (*R <* 10^−19^). (C) When the the rate of environmental change is slightly faster, the prey population cross the extinction threshold (*R <* 10^−19^). At this point, the system is prone to collapse in front of any seemingly negligible environmental noise and demographic perturbation.

**Figure 7:**
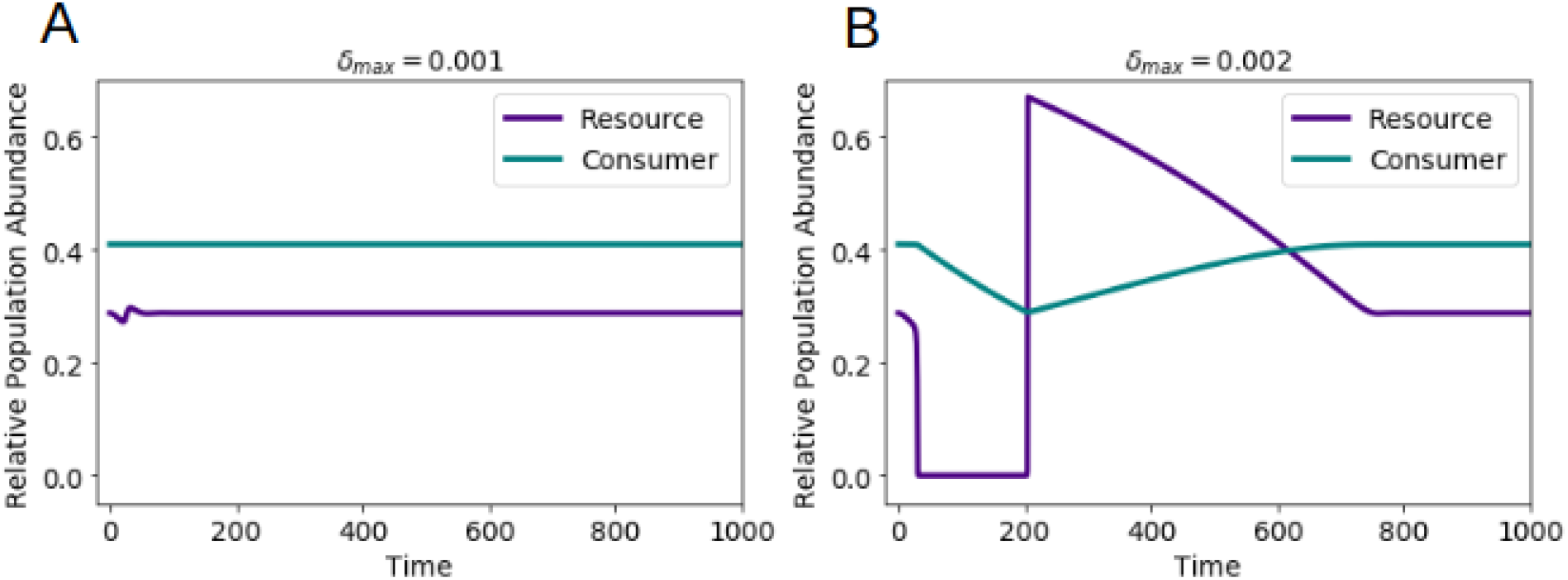
Effect of Magnitude of Environmental Degradation on the system’s vulnerability to Rate-Induced Collapse. Considering the same rate of environmental degradation 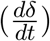 and predator responsiveness (*ϵ*), we tested how different maximal magnitudes of change (*δ*_*max*_) influence system’s susceptibility to rate-induced collapse. For very small and slightly different maximal magnitudes of environmental change (*δ*_*max*_) the system take remarkably distinct trajectories. (A) When the magnitude of environment is smaller (*δ*_*max*_) the prey population never cross the extinction threshold (*R <* 1^−20^). (B) For a slightly larger magnitude of environmental change (*δ*_*critical*_), the prey population cross the extinction threshold (*R <* 1^−20^). At this point, the system is prone to collapse in front of any seemingly negligible environmental noise and demographic perturbation. After the prey population reaches this critically low abundance the population increases. This happens because there is no loss in stability of the coexistence fixed point, hence, the system eventually converges to the equilibrium where both population coexists.

